# Tumor suppressor p53 deficiency increases tumor immunogenicity through promoting IL33-mediated anti-tumor immune responses

**DOI:** 10.1101/2022.09.11.507505

**Authors:** Yang Li, David Shihong Gao, Lixian Yi, Fei Gao, Runzi Sun, Kevin Kai Lu, Junchi Xu, Jason Shoush, Zoi Kykrou, Minxin Liang, Binfeng Lu

## Abstract

Recent studies have shown that p53 contributes to poor survival during immune checkpoint blockade (ICB) therapy. Lung cancer patients with p53 mutations have significantly improved response rates to PD-1 ICB therapy. While previous studies have shown that tumor-derived IL-33 is required for the anti-tumor immune response and efficacy of ICB therapies, the relationship between p53 and IL-33 during ICB therapy is unknown. In this study, we characterized the role of the p53/IL-33 axis in regulating the tumor microenvironment (TME) in response to ICB therapy. CRISPR-Cas9-mediated deletion of Trp53 in tumor cells combined with PD-1 ICB therapy synergistically inhibited tumor growth in a murine MC38 colon adenocarcinoma model. We observed increased CD4+ and CD8+ T cell infiltration, as well as reduced Treg infiltration. IL-33 was upregulated and its expression increased with time and response to treatment. Simultaneous deletion of Il33 in the MC38 tumor cells reversed the efficacy of PD-1 ICB therapy. ST2^-/-^(IL-33 receptor) mice with Trp53-deficient MC38 tumors also showed no response to PD-1 ICB. Our findings depict a novel mechanism by which the loss of p53 in tumors treated with ICB therapy induces upregulation of tumoral IL-33 and host ST2 signaling. p53 mutations may be a double-edged sword for cancer, i.e. loss of the tumor suppressor initially facilitates tumorigenesis, but also leads to upregulation of danger signals in the tumor. These danger signals, such as IL-33, mediate the anti-tumor effect of ICB.

## Introduction

p53 is a well-studied tumor suppressor gene. However, its role in immunotherapy is not clear. p53 mutations have been found to be associated with increased lymphocytes in a large scale pan-cancer genomics and transcriptomic study (*1*). In human non-small cell lung cancer (NSCLC) patients, p53 mutations significantly associated with levels of immune checkpoint molecules, activated T-effector genes, and IFNG signature genes (*2, 3*). NSCLC patients with co-occurring TP53/KRAS mutations showed improved clinical benefit after treatment with PD-1 inhibitors (*3*). In lung cancer patients treated with PD-1 inhibitors, p53 mutations have been associated with better overall response rate and overall survival (*4-6*). For relapsed/refractory (R/R) acute myeloid leukemia (AML) patients treated with Flotetuzumab, an investigational CD123xCD3 bispecific dual-affinity retargeting antibody (DART) molecule, higher expression of immune markers such as IFNG, FOXP3, and immune checkpoints was observed in primary bone marrow samples with p53 mutations when compared to those with wild-type p53. In addition, p53 mutations and deletions are associated with clinical response to Flotetuzumab immunotherapy in AML (*7*). It has been suspected that the increased immunity against p53 mutated cancers is due to increased mutations where p53 function is compromised. Nonetheless, the exact mechanism of how p53 mutation promotes immune responses is not understood.

IL-33 is either constitutively or inducibly expressed in epithelial cells of lining tissues such as the lung, skin, GI tract, breast, and pancreas. Carcinomas are derived from epithelial cells. Compared to normal tissues, IL-33 is downregulated in many carcinomas, such as human lung cancer cells, breast cancer, and cervical cancers. In addition, IL-33 is downregulated in more advanced cancers compared to early stage cancers (reviewed in (*8, 9*). These results suggest that IL-33 downregulation is associated with malignancy. The genetic program that mediates IL-33 downregulation in cancer is not well understood.

In order to study the role of p53 in immunotherapy, we deleted p53 in mouse tumor cell lines. We determined the effect of p53 deletion in PD-1-mediated immunotherapy in mice models. We focused on whether the antitumor activity of IL-33 is regulated by p53.

## Results

### p53 in tumor cells blocks response to cancer immunotherapy

We definitively tested whether p53 affects the efficacy of checkpoint blockade immunotherapy by using a loss of function approach in the transplant mouse model of MC38 colon adenocarcinoma. Using CRISPR-Cas9 in MC38 tumor cells *in vitro*, we targeted for deletion a 70 nucleotide region of exon 4 in the *Trp53* gene that was located in the DNA-binding domain (Figure 1A). Deletion of this region drastically reduced *Trp53* mRNA expression *in vivo* (Figure 1B). In *in vitro* functional assays, where DNA damage was induced by 5-Fu, the p53-deficient cells were unable to upregulate p21 – a cyclin-dependent kinase inhibitor that halts the cell cycle and is a key component of the DNA damage response – as opposed to unedited control or WT cells (Figure 1C). These data show that we have generated a Trp53-null MC38 cell line, which we hereafter term Trp53^KO^ cells, along with their unedited controls, Trp53^CON^ cells.

**Figure 1.**
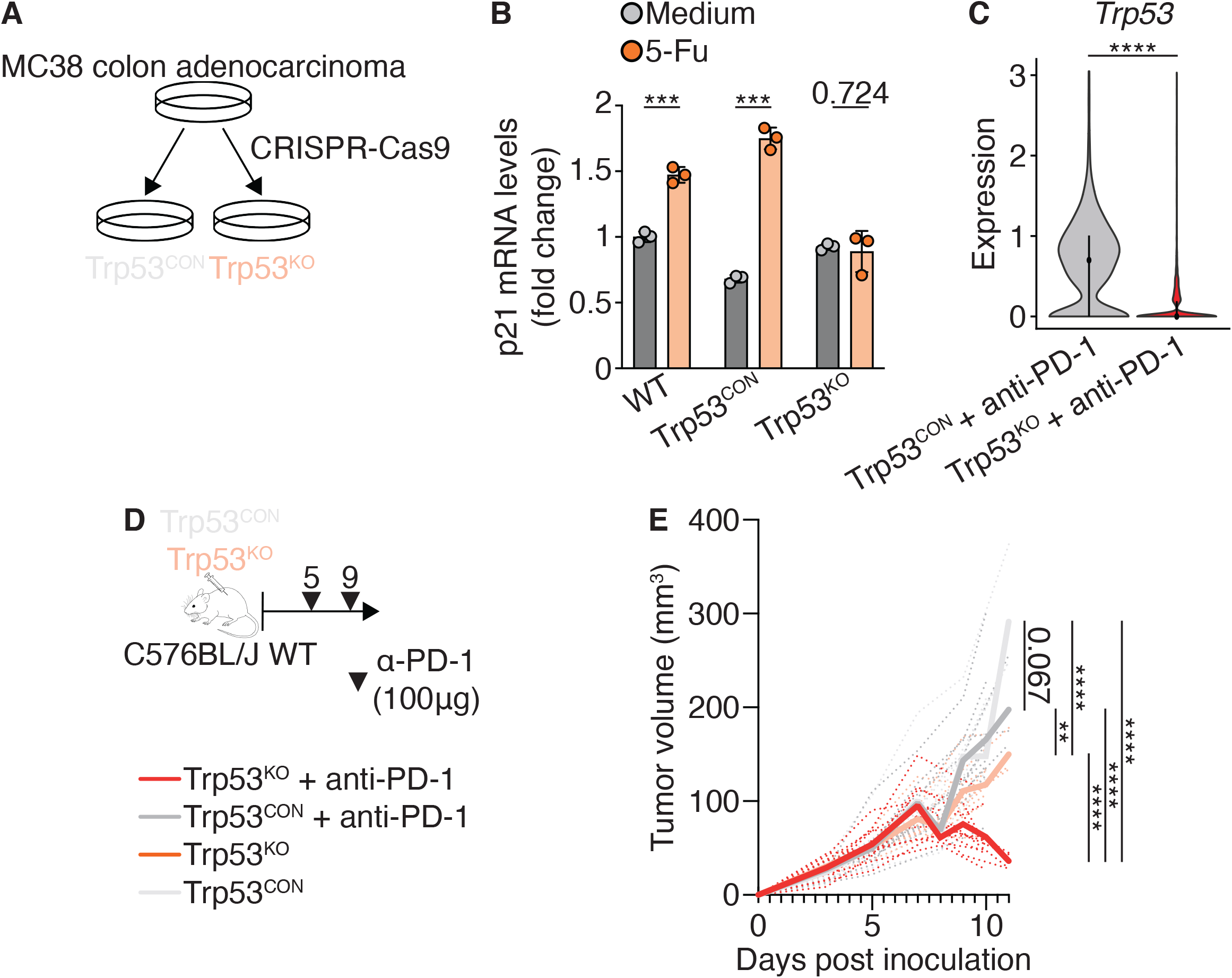
p53-deficiency in tumor cells increases response to cancer immunotherapy. a)Genetic knockout of p53 from MC38 tumor cells using CRISPR-Cas9 to generate p53^CON^ and p53^KO^ tumor cells. b)MC38 WT, p53^CON^, and p53^KO^ tumor cells were treated with medium or 5-Fu, and p21 mRNA levels were measured by qPCR. c)p53 expression in p53^CON^ and p53^KO^ tumor cells from tumors treated with anti-PD-1 therapy by scRNAseq (Figure 2). d-e) p53^CON^ and p53^KO^ tumor cells were inoculated into C57BL/6J mice, treated with anti-PD-1 therapy, and tumor growth measured.

We next tested the response of Trp53^CON^ and Trp53^KO^ cells to anti-PD-1 therapy *in vivo* (Figure 1D). As early as day 9 – a timepoint at which anti-PD-1 therapy does not yet affect the growth of WT tumors – treated Trp53^KO^ tumors already had a significantly smaller volume compared to both untreated and treated Trp53^CON^ tumors. These differences remained at later timepoints, with 1/3 of tumors undergoing complete remission (Figure 4G). Interestingly, untreated Trp53^KO^ tumors also grew slower than untreated and treated Trp53^CON^ tumors, though at a faster rate than their treated counterparts. Our results provide functional evidence that p53 inhibits the efficacy of cancer immunotherapy.

### p53 deficiency does not induce tumor cell-intrinsic changes, but alters the immune response

We next determined whether tumor cell-intrinsic or -extrinsic immune response changes were responsible for the reduction in tumor growth. Trp53^CON^ and Trp53^KO^ cells were inoculated into immunodeficient RAG1^-/-^ (lack B and T cells) and NSG (lack B, T, and NK cells) mice, along with WT controls (Figure 2A). We found no difference in tumor growth between the Trp53^CON^ and Trp53^KO^ cells in either immunodeficient mice strains (Figure 2B-C). These data show that alterations to the immune response, and not cell-intrinsic changes, induced by p53 deficiency in tumor cells, are required for the reduction of tumor growth and improved efficacy of checkpoint immunotherapy.

**Figure 2.**
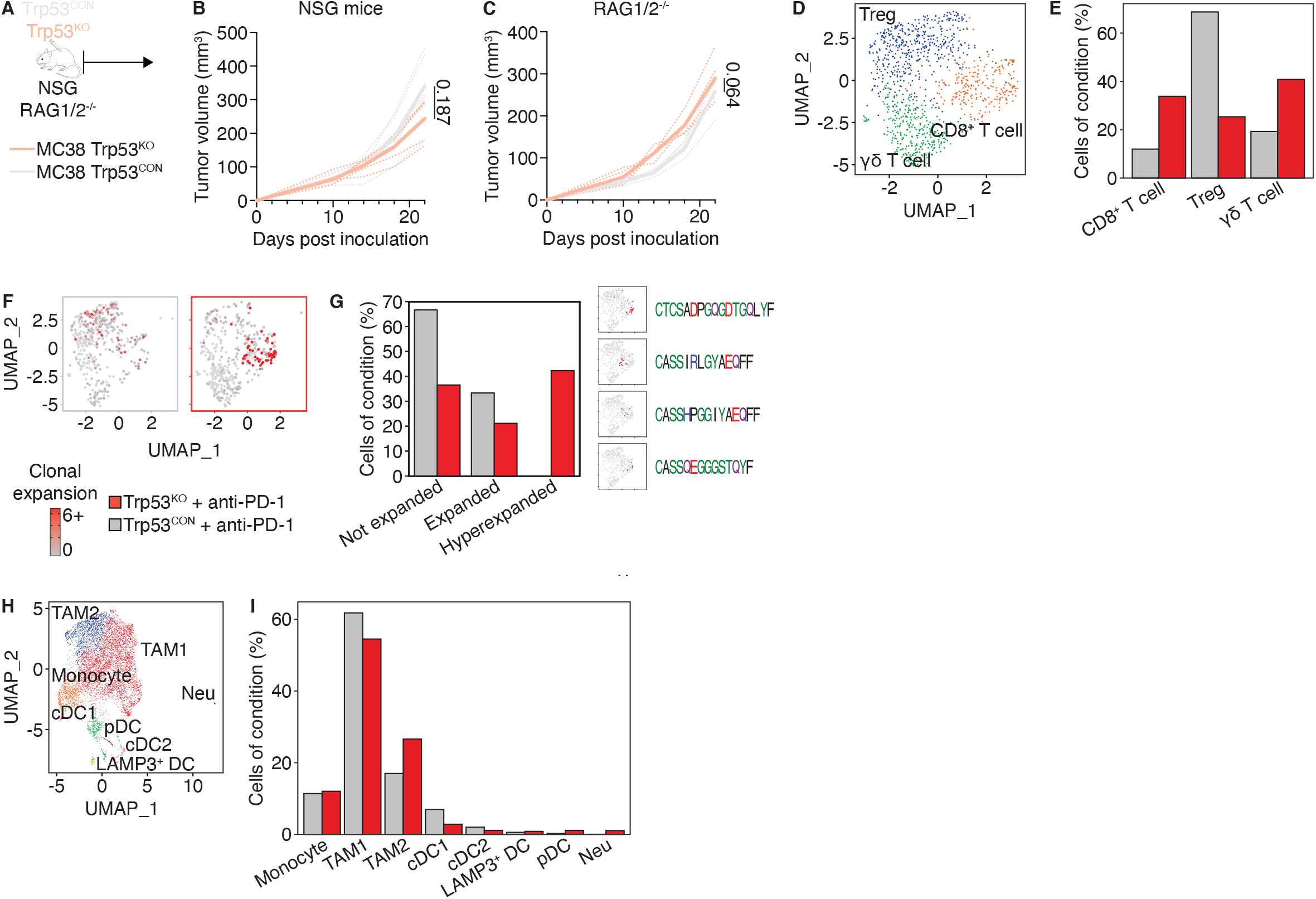
p53-deficient tumor cells do not undergo intrinsic changes, but activate clonal T cell responses a-c) p53^CON^ and p53^KO^ tumor cells were inoculated into RAG1^-/-^ and NSG mice, treated with anti-PD-1 therapy, and tumor growth measured. d-i) Paired scRNAseq and scTCRseq was performed on day 10 tumors generated as in Figure 1D. d)UMAP representation of T cells. e)Changes in T cell populations by condition. f)Clonal expansion of T cells by condition. g)Not expanded (0 cells), expanded (1-5), and hyperexpanded (>5) CD8^+^ T cell clones by condition. Hyperexpanded clones depicted on UMAP along with TCR amino acid sequence. h)UMAP representation of myeloid cells. i)Changes in myeloid cell populations by condition.

### Characterization of the immune response induced by p53-deficient tumor cells during anti-PD-1 therapy

We next characterized how p53 deficiency in tumor cells alters the immune response. We performed single cell RNA and TCR sequencing (scRNAseq and scTCRseq) of anti-PD-1 treated Trp53^CON^ and Trp53^KO^ tumors on day 11 (Figure 2D-I). scRNAseq data showed that CD8^+^ T cells and γδ T cells increased in Trp53^KO^ tumors, while Treg cells decreased (Figure 2E). While CD8^+^ T cells in Trp53^CON^ tumors had modest clonal expansion and clonal size, cells in Trp53^KO^ tumors had hyper-expansion of a few clones (Figure 2F-G). This expansion may be due to neoantigens induced by the loss of p53 and resulting genome instability. Interestingly, the TCR amino acid sequences of these hyper-expanded clones had some commonalities, such as a starting CASS motif, glycine-rich middle, and ending Q and F residues (Figure 2G). Treg cells, which have a lower specificity and affinity for the MHC-peptide complex, had a smaller clonal size, but showed widespread expansion in Trp53^CON^ tumors; this expansion was drastically reduced in Trp53^KO^ tumors (Figure 2F-G). In the myeloid compartment, we did not observe changes to the total monocyte, TAM, or DC populations (Figure 2H-I). However, the phenotype of macrophages in Trp53^KO^ tumors shifted from type 1 to type 2 (Figure 2H-I). In addition, both the cDC1 and cDC2 populations decreased, while LAMP3^+^ DCs and pDCs increased (Figure 2H-I). We also observed an increase in neutrophils (Figure 2H-I).

We validated our characterization of the immune response by performing multicolor FLOW cytometry (Figure 3A). We found that treated Trp53^KO^ tumors had increased CD8^+^ T cells that expressed an intense type 1 signature of IFNG and GZMB, compared to both untreated and treated Trp53^CON^ tumors (Figure 3B). In addition, we also observed an increase in T^conv^ cells and decrease in Tregs (Figure 3C-D). These findings are consistent with our transcriptional data. The total TAM population decreased in treated Trp53^KO^ tumors, but we still found a type 1 to type 2 shift (Figure 3E). The total DC population trended towards an increase, and we again found a decrease in both cDC1 and cDC2s (Figure 3F). We also found a striking decrease in gMDSCs and increase moMDCSs (Figure 3G). Monocytes also decreased and neutrophils increased (Figure 3H-I). Interestingly, we found that untreated Trp53^KO^ tumors showed similar trends to their treated counterparts, albeit at lesser degrees. Our profiling of the immune response consistently showed that p53 deficiency in tumor cells and anti-PD-1 therapy drastically alters the immune response *in vivo*, primarily augmenting T cell responses.

**Figure 3.**
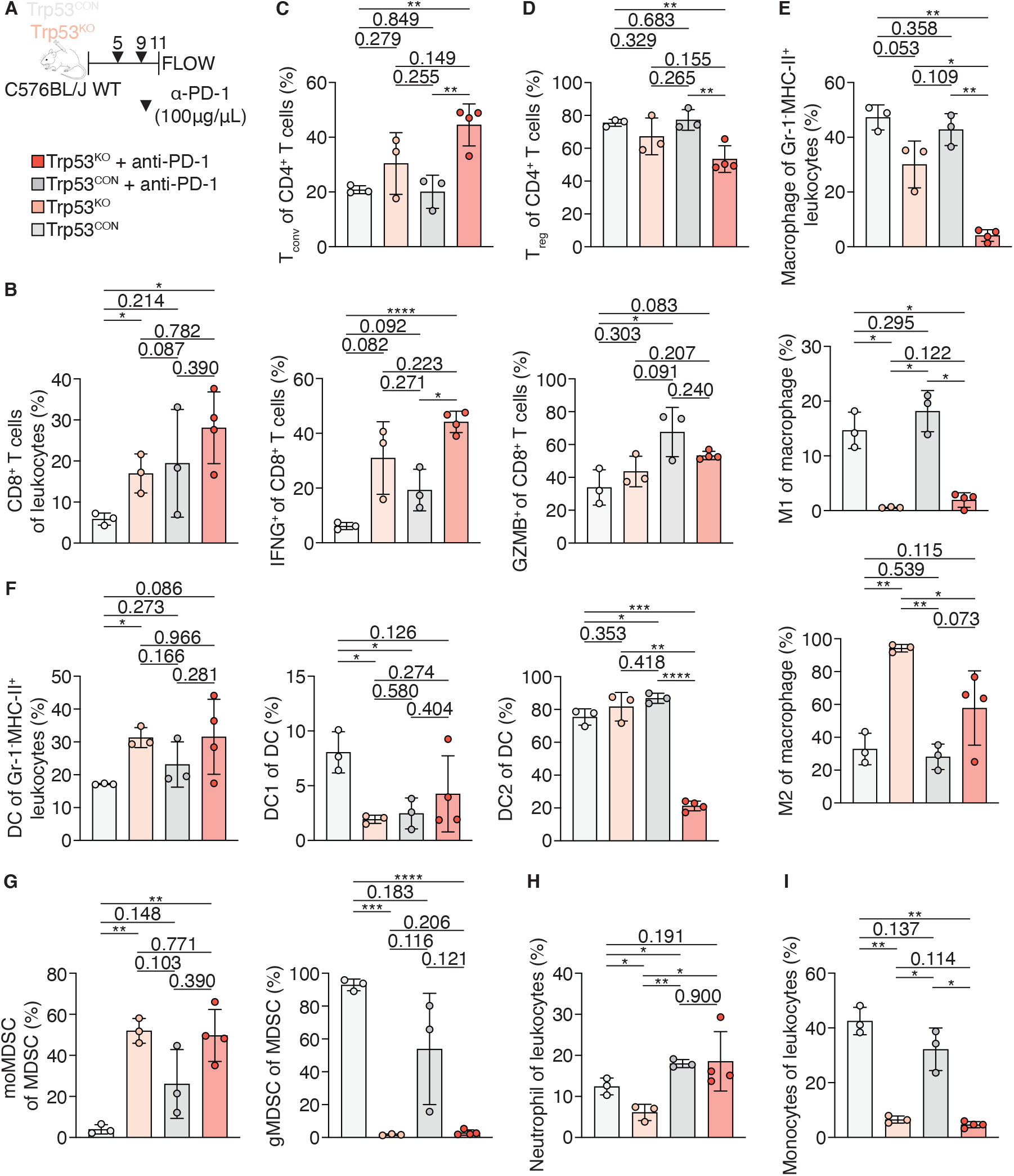
Loss of p53 in tumor cells remodels the immune compartment during anti-PD-1 therapy. a-l) Flow cytometry for immune cell populations was performed on day 10 tumors generated as in Figure 1D.

**Figure 4.**
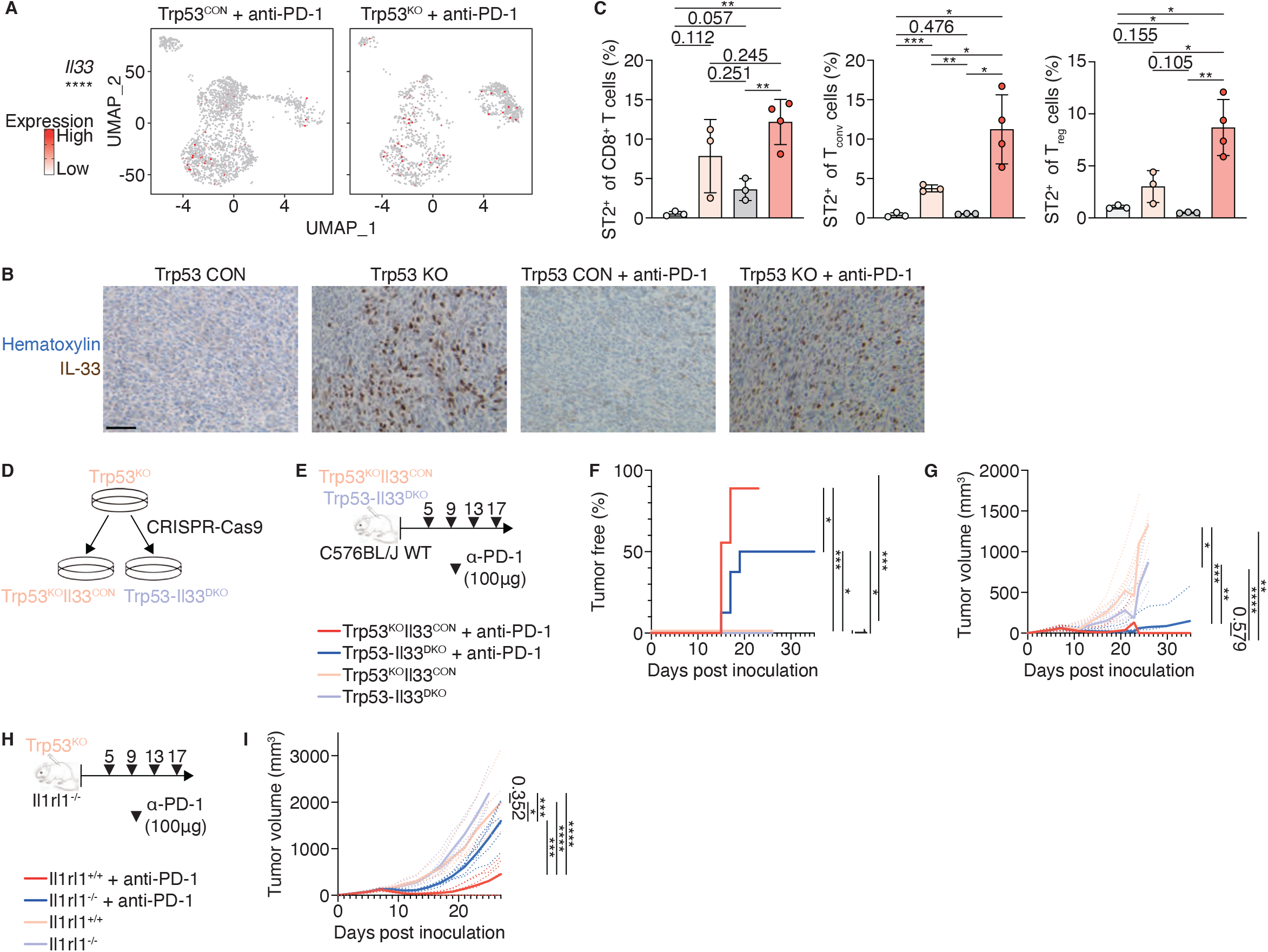
p53 represses IL-33 expression to reduce anti-PD-1 therapeutic efficacy a)Il33 expression in p53^CON^ and p53^KO^ tumor cells from tumors treated with anti-PD-1 therapy by scRNAseq (Figure 2). b)ST2 expression in CD8^+^, T^conv^, and Treg populations as measured by FLOW cytometry as in Figure 3. c)H&E and immunohistochemical staining for IL-33 was performed on day 11 tumors generated as in Figure 1D. d)Genetic knockout of Il33 from MC38 p53^KO^ tumor cells using CRISPR-Cas9 to generate p53^KO^-IL-33^CON^ and p53^KO^-IL-33^CON^ tumor cells. e-g) MC38 p53^KO^-IL-33^CON^ and p53^KO^-IL-33^CON^ tumor cells were inoculated into C57BL/6J mice, treated with anti-PD-1 therapy, and tumor growth measured. h-i) MC38 p53^CON^ and p53^KO^ tumor cells were inoculated into Il1rl1^+/+^ and Il1rl1^-/-^ mice, treated with anti-PD-1 therapy, and tumor growth measured.

### p53 represses IL-33 expression in tumor cells to lower anti-PD-1 therapeutic efficacy

In our scRNAseq data, Trp53^KO^ tumor cells significantly upregulated the alarmin IL-33 (Figure 4A). Immunohistochemistry (IHC) staining for the IL-33 revealed an even more striking increase in the levels of nuclear IL-33 protein (Figure 4B). IL-33 stimulates type 1 and type 2 immunity, as well as Treg cells. Thus, we checked expression of the IL-33 receptor (IL1RL1/ST2) in the immune compartment using our scRNAseq data and flow cytometry (Figure 4C). IL1RL1 was significantly upregulated on CD8^+^ T cells, T^conv^ cells, and Treg cells – all T cells – at both the mRNA and protein levels. These data show that p53 indirectly represses IL-33 expression in tumor cells, and may thereby alter the immune response and tumor growth.

We next tested the effects of p53 regulation of IL-33 by again using a loss of function approach. We used CRISPR-Cas9 to delete a coding region of IL-33 from Trp53^KO^ tumor cells (double knockout [DKO]), thus generating Trp53-Il33^DKO^ cells and unedited Trp53^CON^ Il33^DKO^ cells (Figure 4D). While DKO did not significantly affect tumor growth following anti-PD-1 therapy, it significantly decreased the frequency and rate of complete remissions overtime (Figure 4E-F). We further tested whether IL-33 signaling through IL1RL1 on host cells, which include IL1RL1-expressing T cells, also reverses our phenotype. anti-PD-1 therapy drastically reduced tumor growth of Trp53^KO^ cells in WT mice, but not IL1RL1^-/-^ mice, where growth was closer to the level of untreated tumors (Figure 4G). In both experiments, tumor growth and remission were unaffected without anti-PD-1 therapy, likely due to interplay between the p53-IL-33 axis and the immune response, and indicating that the axis is relevant in the context of immunotherapy. Taken together, these data show that p53 indirectly represses IL-33 expression in tumor cells and thereby reduces response to anti-PD-1 therapy, likely by blocking IL-33 signaling that activates immune cells.

## Discussion

Here, using two independent murine tumor models, we demonstrated that p53 deletion results in an increase of IL-33 expression in tumor cells. p53 deletion decreased tumor growth in immune-sufficient mice, but not in immune compromised mice. In addition, p53 deletion improved the antitumor efficacy of PD-1 blockade tumor immunotherapy in mice. Tumor cell deletion of IL-33 and host mice deficiency in Il1rl1 decreased the antitumor effect of p53 deficiency. These data illustrate that p53 suppresses IL-33 expression and thereby inhibits the immunogenicity of cancer cells.

Many stress and inflammatory signals can induce IL-33 expression. TNF-α and IL-1β synergistically induce IL-33 in skin fibroblasts. IFN-γ upregulates IL-33 in keratinocytes PMID: 23362867. IL-4 and IL-13 increase the expression of full-length IL-33 in the nucleus of keratinocytes (*10*). Double stranded RNA (dsRNA) can enhance IL-33 promoter activity through the TLR3-EGFR-IRF3 pathway in normal human epidermal keratinocytes (*11*). Hypoxia increases IL-33 and ST2 expression by human pulmonary arterial endothelial cells (HPAECs). The IL-33 further boosts hypoxia-induced expression of VEGFA, VEGFR-2, and ICAM-1 and vascular remodeling in hypoxic pulmonary hypertension (*12*). We have also found that immune checkpoint inhibitors treatment can also result in increases of IL-33 expression in mouse tumor cell lines in vivo. The current work has confirmed this finding, further substantiate the link between greater tumor immunogenicity and IL-33 expression in TME.

Many transcription factors either promote or suppress the IL-33 gene expression in a wide range of cell types such as epithelial cells, fibroblasts, endothelial cells, and macrophages. TBX2, a transcriptional repressor that plays an important role in maintaining the proliferation of mesenchymal progenitors during lung development, inhibits IL-33 expression (*13*). Ovol1, a skin disease‒linked transcriptional repressor, binds iL-33 promoter and represses IL-33 expression in skin epithelial cells. And the Ovol1/IL-33 link regulates skin inflammation (*14*). An OCT-1 (POU2F1) binding site overlaps the asthma-associated single nucleotide polymorphism (SNP) rs1888909, which resides within 5kb of IL-33 gene and is associated with IL-33 expression (*15*). Focal adhesion kinase (FAK) can also translocate to the nucleus, where it binds to and promotes IL-33 expression in squamous cell carcinoma (SCC) cell lines (*16*). NFE2L3 can bind the regulatory sequences of the Il-33 locus and promote its expression (*17*) in colorectal cancer cells. CDX2 directly binds and promote the intronic enhancer activity in the IL33 gene in primary colonic epithelial cells from healthy humans and epithelial cell lines (*18*). Friend leukemia virus integration 1 (Fli1), whose deficiency is a predisposing factor of systemic sclerosis (SSc), occupies the IL33 promoter in dermal fibroblasts and represses IL33 expression. Aryl hydrocarbon receptor (AhR) binds to a dioxin response element (DRE) on the IL-33 promoter and activate IL-33 expression in human macrophage cell line THP1 and bronchial epithelial cells

(*19*) (*20*). These published findings highlight the complex regulation of IL-33 in different types of cells. The possible association between p53 and IL-33 has been indirectly shown in a clinical study on relapsed/refractory (R/R) acute myeloid leukemia (AML) patients treated with flotetuzumab. This study shows that IL33 was more highly expressed in TP53-mutated compared with TP53-WT patients (*7*). The mechanism underlying the p53/IL33 axis is likely indirect as shown in this study.

These results are consistent with clinical data demonstrating that p53 mutations are associated with better clinical outcome in ICI therapy for NSCLC patients, (*2-6*) and for R/R AML patients treated with Flotetuzumab (*7*). Here, we provide a potential mechanism where p53 deletion leads to increased danger signals, leading to increase immunogenicity and better responses to immunotherapy.

## Materials and methods

### Mice

Mice were maintained in a specific-pathogen-free animal facility in accordance with an animal protocol approved by the Institutional Animal Care and Use Committee of the University of Pittsburgh. WT (stock #000664) and RAG1^-/-^ (B6.129S7-*Rag1*^*tm1Mom*^/J, 002216) mice on the C57Bl/6J background, as well as NSG (NOD.Cg-*Prkdc*^*scid*^ *Il2rg*^*tm1Wjl*^/SzJ, 005557) were purchased from the Jackson laboratory. *Il1rl1-deficient* mice on the C57BL/6 background were kindly provided by Dr. Andrew McKenzie (MRC Laboratory of Molecular Biology) and maintained in our facility (*21*).

### Tissue culture

MC38 colon adenocarcinoma cells were maintained in Dubecco”s Modified Eagle Medium, high glucose supplemented with 10% fetal bovine serum and 1% penicillin-streptomycin.

### Knockout of Trp53 in MC38 cells

Trp53 and Il33 knockout were performed using CRISPR-Cas9. Trp53;

gRNA1: ACAGCCATCACCTCACTGCA, gRNA2: ACACTCGGAGGGCTTCACTT, gRNA3:

GTGACAGGGTCCTGTGCTGC, gRNA4: ACAGGGGCCATGGAGTGGCT. Il33 knockout was performed as previously described (*22*).

### Tumor models

Mice were shaved and inoculated intradermally with Trp53^CON^ and Trp53^KO^ MC38 cells at 10^6^ cells/50uL PBS/tumor. One tumor was inoculated per mouse. Tumor length (L) and width (W) were measured using a digital caliper. Tumor volume was calculated using the formula: L x W x W x 0.5. Tumors were treated with anti-PD-1 purchased from BioxCell *InVivo*MAb anti-mouse PD-1 (catalog #BE0146). Antibodies were aliquoted and diluted in PBS to the required concentration to inject each mice intraperitoneally with 100ug in 100uL. Mice were treated on days 5, 9, 13, and 17, and tumors were allowed to grow up beyond the last treatment.

### Tissue processing

Mice were euthanized and tumors immediately dissected. Excess skin, hair, fat, and connective tissue were removed. Tumors were transferred into 6-well plates containing 0.25mg/mL LiberaseTL (Roche, 5401020001) and 0.33mg/mL DNase (Sigma-Aldrich DN25-10MG) in RPMI, minced with scissors, and digested in a tissue culture incubator for 30min without agitation.

Digestion was quenched with 3mL RPMI. Samples were strained through 70-uM strainers and pushed through using a pestle, then strained through 30uM nylon mesh. Samples were pelleted and resuspended in 2% FBS in HANKS.

### Flow cytometry

Staining was performed in 96-well V-bottom plates. For cytokine analysis, cells were first stimulated using the Leukocyte Activation Cocktail, with BD GolgiPlug (BD Biosciences 550583) in a tissue culture incubator for 4 hours. Cells were initially stained in 0.1% Ghost Dye Violet 510 (Tonbo Biosciences, 13-0870-T500) in PBS on ice for 30min. They were then stained in antibody cocktails in 2% FBS in HANKS on ice for 5 min. Cells were then filtered through 30uM nylon mesh. Samples were run on a Cytek Aurora.

The following antibodies were used: ArgI (eBioscience, catalog #46-3697-82, clone A1exF5), CD4 (BD Biosciences, 612844, RM4-5), CD8a (BD Biosciences, 566096, 53-6.7), CD11b (BD Biosciences, 564443, M1/70), CD11c (Biolegend, 117312, N418), CD24, CD45 (BioLegend, 103130, c30-F11), CD44, CD62L, CD80, CD86 (BioLegend, 105043, GL-1), CD206 (BioLegend, 141708, C068C2), F4/80 (BioLegend, 123130, BM8), Gr-1, Ly-6C (BioLegend, 128037, HK1.4), Ly-6G (BioLegend, 127628, 1A8), MHCII, PD-1 (BD Biosciences, 744633, J43), Tim-3 (BioLegend, 119721, RMT3-23), LAG-3 (BioLegend, 125212, C9B7W), ST2 (eBioscience, 46-9333-82, RMST2-33), CD103 (BD Biosciences, 565849, M290), TCF1 (Cell Signaling Technology, 14456S, C63D9), Foxp3 (BioLegend, 126410, MF-14), GzmB (BioLegend, 372204, QA16A02), Ki-67 (BioLegend, 652410, 16A8), and IFN-γ (BioLegend, 505826, clone XMG1.2). All antibodies were used at 1:100 dilution.

### scRNAseq

Dissected and trimmed tumors were washed in pre-cooled RNase-free H^2^O and transferred into MACS Tissue storage solution (Miltenyi Biotec, 130-100-008). Samples were shipped on ice overnight to Novogene. Samples were processed according to the standard 10X pipeline. Three tumors from separate mice were prepared separately for each condition. The sample with the highest best cell viability and concentration for each condition was chosen. From the two samples, 10000 cells (target 7000 cells in output) per sample were loaded onto two separate lanes of a Chromium Chip in a Chromium Controller (10X Genomics) with 5” chemistry for gene expression and TCR analysis. Libraries were sequenced on an Illumina Novaseq 6000 S4 with 30000 reads/cell (∼200M reads total, 60Gb data output).

Cellranger 2 was used to align fastq files were aligned to the mm10 reference genome and generate barcode, feature, and count matrices. Initial downstream analysis was performed using Seurat. Custom code was used for further analyses.

### Immunohistochemistry

Immunohistochemistry was performed as previously described in ((*23*)) using IL-33 antibody (R&D, AF3626).

## Data availability

Data generated is available upon request to the corresponding author.

## Contributions

Conceptualization: B.L., Y.L., D.S.G. Experimentation and analysis: Y.L., D.S.G., L.Y., F.G., R.S., K.K.L., J.X., J.S., Z.K., M.K. Computational analysis: D.S.G. Writing, original draft: D.S.G. Writing, review and editing: D.S.G. and B.L. Supervision: B.L.

